# The GridCAT: A toolbox for automated analysis of human grid cell codes in fMRI

**DOI:** 10.1101/106096

**Authors:** Matthias Stangl, Jonathan Shine, Thomas Wolbers

**Author notes:** These authors contributed equally to this work (Matthias Stangl, Jonathan Shine). Corresponding Author: Matthias Stangl German Center for Neurodegenerative Diseases (DZNE) Leipziger Str. 44, 39120 Magdeburg, Germany.

## Abstract

Human fMRI studies examining the putative firing of grid cells (i.e., the grid code) suggest that this cellular mechanism supports not only spatial navigation, but also more abstract cognitive processes. This research area, however, remains relatively unexplored, perhaps us to the complexities of data analysis. To overcome this, we have developed the Matlab-based Grid Code Analysis Toolbox (GridCAT), providing a graphical user interface, and open-source code, for the analysis of fMRI data. The GridCAT performs all analyses, from estimation and fitting of the grid code in the general linear model, to the generation of grid code metrics and plots. Moreover, it is flexible in allowing the specification of bespoke analysis pipelines; example data are provided to demonstrate the GridCAT’s main functionality. We believe the GridCAT is essential to opening this research area to the imaging community, and helping to elucidate the role of human grid codes in higher-order cognitive processes.

**Highlights:**

- The putative firing of grid cells (i.e., the grid code) can be examined using fMRI
- Necessary steps for grid code analysis are reviewed
- The Matlab-based grid code analysis toolbox (GridCAT) is introduced
- Automated grid code analysis can be conducted either via a graphical user interface or open-source code
- A detailed manual and an example dataset are provided

## 1. Introduction

Identifying the neural mechanisms supporting spatial navigation remains a key goal for neuroscience. In the last 10 years, significant progress has been made with the discovery of the grid cell in the rat medial entorhinal cortex, a neuron exhibiting firing properties that could provide a spatial metric underlying navigational functions such as path integration (Hafting et al., 2005). Grid cells have been found subsequently in a diverse range of mammalian species (for a detailed review, see Rowland et al., 2016), and, more recently, the putative signature of grid cell firing, which we refer to as the grid code throughout this paper, has been identified also in healthy human subjects using fMRI (functional magnetic resonance imaging; Constantinescu et al., 2016; Doeller et al., 2010; Horner et al., 2016; Kunz et al., 2015). Given the increasing interest in the role of grid cells in human cognition, and the absence of standard analysis tools to examine grid codes in fMRI, we present here a Matlab-based toolbox, the Grid Code Analysis Toolbox (GridCAT), which generates grid code metrics from functional neuroimaging data. The GridCAT is openly available at http://www.wolberslab.net/gridcat

Unlike place cells that have single firing fields (O’Keefe & Dostrovsky, 1971), grid cells in the rat medial entorhinal cortex fire in multiple different locations within the environment (Fyhn et al., 2004; Hafting et al., 2005). The firing fields of these cells show remarkably regular organisation, forming tessellating equilateral triangles that effectively ‘tile’ the world’s navigable surface in a hexagonal lattice (Hafting et al., 2005). The equally spaced and repetitive firing means that, for each firing field of a grid cell, the six adjacent fields are arranged in 60^o^ intervals, creating a six-fold symmetry. Each cell’s grid can differ, however, in several ways, such as its orientation, spacing, and the size of its firing field. Although recording from a population of grid cells within the entorhinal cortex reveals variability in these properties, there appears to be topographical arrangement of these cells. For example, the grids of neighbouring cells are more similar in terms of their orientation and spatial scale relative to cells located further apart (Hafting et al., 2005); distal cells, however, can still show coherence in the orientation of their grids, even though their spatial scales may differ (Barry et al., 2007). Given that each cell’s grid is spatially offset (to varying degrees) relative to a neighbouring one, it has been hypothesised that these cells may provide the neural mechanism for complex spatial navigation abilities such as path integration (Hafting et al., 2005).

Grid cells have been found in a number of other species, including bats (Yartsev et al, 2011), primates (Killian et al., 2012), and humans (Jacobs et al., 2013). Although rare and often comprising small sample sizes, studies using intracranial recordings in humans provide an opportunity to use experimental methods analogous to those routinely used in behavioural neuroscience (i.e., recording directly from neurons). Jacobs et al. (2013) recorded from cells in the entorhinal cortex of patients with intractable epilepsy as they completed an object-place memory task requiring them to navigate a virtual environment. Consistent with the rat electrophysiology, there was evidence of cells with a six-fold symmetry in their firing rate. The grid cell, therefore, appears preserved across different mammalian species, including humans, and may comprise a common neural mechanism for spatial navigation.

FMRI is used commonly to investigate the neural correlates of higher-order cognitive processes in large samples of healthy subjects, and this method has been applied to the study of putative grid cell firing (Doeller et al., 2010). Although fMRI is able only to detect changes in signal over thousands of neurons, several properties of grid cell firing suggested it would be possible to detect grid codes in the blood oxygenated-level dependent (BOLD) response at the macroscopic level. First, as described earlier, even though grid cells are arranged topographically, the grid orientation of distal cells may still be coherent (Barry et al., 2007). This means that the global signal associated with the firing of these neurons would be consistent across different voxels. Second, it has been shown that the firing of ‘conjunctive’ grid cells in rats is modulated also by travel direction (Sargolini et al., 2006), and their preferred firing orientation is aligned with the main axes of the grid (Doeller et al., 2010). These differences in the dynamics of grid cell firing could be reflected in a six-fold sinusoidal pattern observable in the BOLD response (i.e., the grid code) when participants performed translations either aligned or misaligned with the grid’s axes (Figure 1). Using an object-place memory task in a virtual environment, Doeller et al. (2010) found exactly this pattern of data in several brain regions, including the entorhinal cortex. Consistent with the results of rodent electrophysiology, the BOLD signal showed a six-fold symmetry, with greater activity associated with translations in which the travel direction was aligned with the mean grid orientation, compared to when the travel path was misaligned with a grid axis (the precise methods for estimating the mean grid orientation, and testing the model, are described in detail below). This study, therefore, was critical in demonstrating that fMRI could be used to study grid codes in humans.

**Figure 1.**
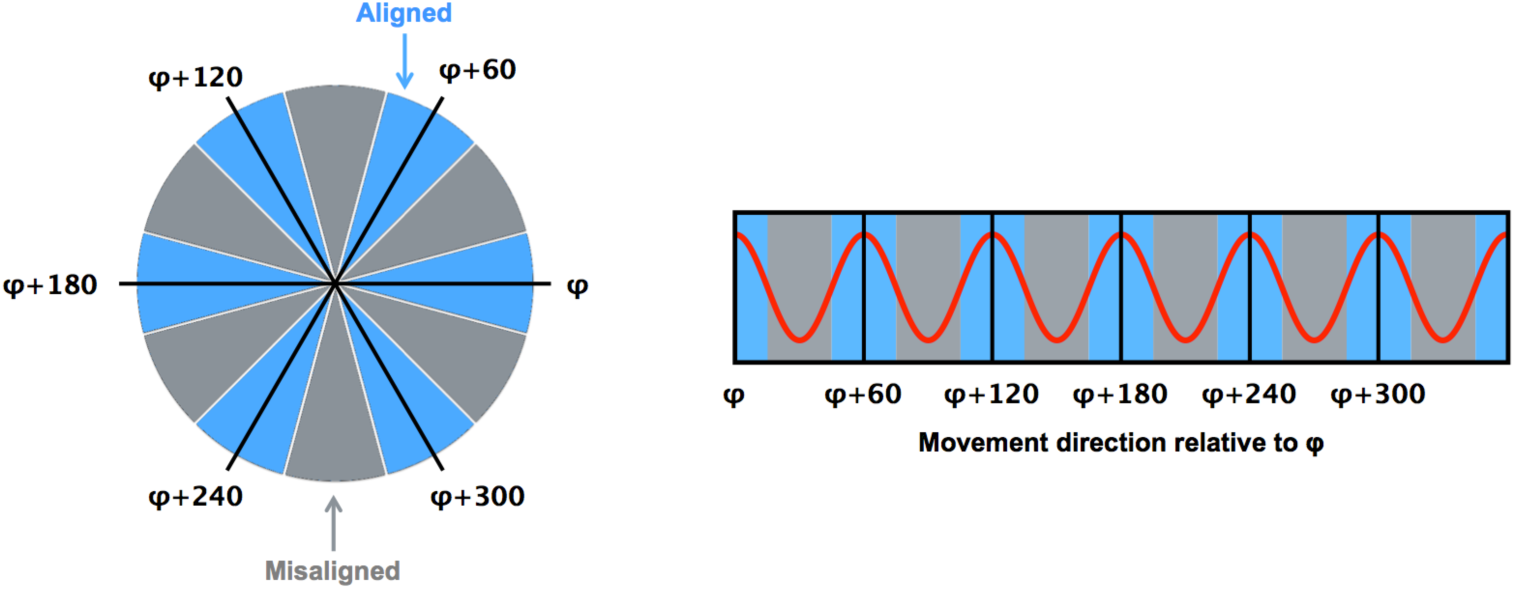
The logic for grid code analysis in human fMRI. Left: Movement directions can be categorized either as aligned (blue) or misaligned (grey) with the mean grid orientation (φ). Right: The red curve shows the expected pattern of the BOLD signal modulated by movement direction relative to φ. Positive signal peaks are expected for movements aligned with φ or a 60 degree multiple of φ (blue sectors). Figure adapted from Doeller et al. (2010).

The identification of grid codes using fMRI has already generated a number of promising new research questions. For example, the estimation of grid codes has been shown to have potential clinical applications with reduced grid cell coherence evident in those at increased genetic risk of Alzheimer’s disease (Kunz et al., 2015). Furthermore, although there appear to be commonalities in the neural mechanisms supporting navigation across diverse species, the study of grid codes in fMRI has demonstrated that these spatial codes may be used more flexibly in humans (Horner et al., 2016). Specifically, Horner et al. found evidence of the sinusoidal pattern in the BOLD response when participants imagined navigation in a virtual environment, despite the absence of visual input. Finally, Constantinescu et al. (2016) demonstrated that recently acquired conceptual knowledge is organised using the same six-fold spatial symmetry. In humans, therefore, these grid codes may be used more abstractly in service of higher-order cognitive processes beyond pure spatial navigation.

Given the recent increase in interest in the function of grid cells in humans, and the distinctly non-trivial method of estimating grid codes in the BOLD signal, the GridCAT allows users to input easily their study design and performs all analyses to estimate the grid code in functional images, using either the graphical user interface (GUI) or the open-source code provided with the toolbox. Furthermore, it requires only a basic Matlab installation and SPM (i.e., no additional Mathwork toolboxes are required), and is compatible with Windows, Linux, and Mac OS. A detailed manual guides users through all steps of a grid code analysis and an example dataset is provided with the GridCAT to explore its functionality. The GridCAT, therefore, opens up the field of grid code research to the wider neuroscience community.

Although there are similarities across fMRI studies in the methods used to estimate grid code metrics, there is as yet no standard analysis pipeline. Because of this, the GridCAT has been designed to be flexible in accommodating a number of different analysis options; decisions regarding a researcher’s own pipeline will depend upon paradigm-specifics and the research question of interest. We note that a recent grid code study examined the neural signal associated with imagined trajectories in the environment (Bellmund et al., 2016). We do not discuss this experiment here, however, because, rather than the mass univariate method commonly used in the study of grid codes, they used multivariate representational similarity analysis, for which there are already several toolboxes available (e.g., Nili et al., 2014; Oosterhof et al., 2016). In the following section, we review extant methods for deriving grid codes in fMRI, and highlight differences in analysis approaches. The aim of this review is to inform the GridCAT user of the different analysis options that have been used previously, and that are available in the toolbox, rather than to provide a critique as to best practice for deriving grid code metrics.

## 2. Grid code analysis

Although analysis pipelines for the examination of grid codes using fMRI differ in several aspects (see Sections 2.1 – 2.6), the overall procedure is relatively similar. First, events of interest for the grid code analysis (i.e., grid events) are specified in the time course of the imaging data. Second, the imaging data are then partitioned into estimation and test datasets. Third, a general linear model (GLM) is fit in the estimation dataset to estimate voxel-wise orientations of the grid code (i.e., GLM1 – see Section 2.4). Fourth, these voxel-wise orientation values are then averaged over voxels in a region of interest (ROI) to generate a mean grid orientation used for a second GLM in the test dataset where grid events are modelled with respect to their alignment with the mean grid orientation (i.e., GLM2 – see Sections 2.6.1 – 2.6.2). Finally, grid code metrics are computed, such as the magnitude of grid code response as well as measures of between-, or within-voxel orientation coherence of the grid code (see Section 2.6.3). In the following sections, we provide more information regarding these individual steps of the grid code analysis pipeline.

### 2.1. Functional image preprocessing for grid code analysis

The GridCAT is agnostic with regards to the nature of the preprocessing carried out on functional images prior to the grid code analysis. For example, the analysis can be conducted using a participant’s normalized, and smoothed, functional images (Constantinescu et al., 2016; Doeller et al., 2010; Horner et al., 2016). Alternatively, one could work in the individual subject’s native functional space (Kunz et al., 2015). Motivations for normalizing to standard space prior to analysis include the desire to examine group-level, cluster-statistics (e.g., Constantinescu et al., 2016), whereas researchers concerned about spatial distortions or interpolation errors in their data resulting from normalization to a standard template might choose to perform the grid code analysis in the participant’s native space.

### 2.2. Specifying grid events

Before the grid code can be estimated, it is necessary to specify grid events within the fMRI time course. For example, grid events could comprise periods of translational movement (e.g., Doeller et al., 2010; Horner et al., 2016; Kunz et al., 2015) within a virtual environment. For each grid event, an angle relative to a nominal 0^o^ reference point (e.g., a fixed landmark in the virtual environment) is then defined, resulting in the ‘grid event angle’.

### 2.3. Partitioning the grid code data into estimation and test sets

Given that the functional data are labelled either as estimation or test data, researchers must decide how to perform this partition. One method is to split the data run-wise into odd and even runs (Doeller et al., 2010; Kunz et al., 2015), performing the estimation in the odd runs and testing in the even ones (or vice-versa). Alternatively, one could split the data into *n* number of temporal bins, and perform the same analysis on these odd/even bins (Horner et al., 2016). As well as offering these data partitioning methods, the GridCAT also provides options to separate grid events within each scanning run into odd and even events, or to split each scanning run into two halves so that estimation and test are calculated on the first and second halves of runs, respectively. Furthermore, if these default partitioning options are not suitable for a particular experiment, bespoke partitioning schemes can be specified in the GridCAT event-table (which is described in detail in the GridCAT manual), allowing the user to specify whether a particular grid event should be assigned either to the estimation or test datasets.

### 2.4. Estimating voxel-wise grid orientations in the BOLD signal

For the estimation data (GLM1), the grid event angle is used to create two parametric regressors for the grid events, using sin(α_t_*6) and cos(α_t_*6), respectively, where α_t_ represents the grid event angle. The multiplication term (*6) used in the calculation of these two regressors transforms the grid event angle into 60^o^ space, mirroring the hexagonal symmetry observed in grid cell firing. By including these parametric regressors in the general linear model, voxels with time courses showing modulation of their signal according to six evenly spaced 60^o^ intervals would have parameter estimates with high absolute amplitudes. The mean grid orientation across all voxels in an ROI can then be calculated. To compute the mean grid orientation, the beta estimates (β1 and β2) associated with the two parametric regressors are each averaged over all voxels in the ROI, and the resulting two values submitted to: arctan[mean(β1)/mean(β2)]/6.

Once the mean grid orientation has been calculated, this value can be used to categorise individual grid event angles in the test data (GLM2) to determine different grid code metrics. For example, grid event angles could be classified either as aligned or misaligned with the mean grid orientation (see Section 2.6).

### 2.5. ROI selection

As described above, the mean grid orientation can be calculated in any chosen ROI, providing that the mask and functional data are registered to one another. Popular choices of ROI include anatomical masks, such as the entorhinal cortex (Doeller et al., 2010; Horner et al., 2016; Kunz et al., 2015), however it is possible also to input to the GridCAT a functionally-defined mask from an orthogonal contrast (e.g., Constantinescu et al., 2016), or localiser dataset.

### 2.6. Quantifying the magnitude of the grid code response

The greatest degree of heterogeneity in analysis pipelines of fMRI grid code studies stems from how the grid code is quantified, or the choice of grid code metric. This relates, in part, to the research question of interest, and we outline here the different metrics that have been used thus far in the published literature. It is worth noting that these methods are not mutually exclusive, and a researcher may want to use a combination of different approaches to test a number of different hypotheses.

#### 2.6.1. Parametric modulation

In the original study reporting grid codes in the fMRI signal, Doeller et al. (2010) fitted a parametric regressor to the grid events in the test data to examine whether voxels in an entorhinal cortex ROI showed evidence of a six-fold sinusoidal pattern of activity. The parametric regressor was calculated by taking each grid event angle (α_t_), and determining its difference from the mean grid orientation (φ) by calculating cos[6*(α_t_ – φ)], which resulted in values ranging between ‘1’, for grid event angles aligned perfectly with the mean grid orientation (or a 60^o^ multiple of it), and ‘-1’ for values completely misaligned with the grid code phase (i.e., mean grid orientation + 30^o^, plus any 60^o^ multiple of this value). Using cluster statistics, Doeller et al. reported voxels at the group-level showing modulation of their signal according to this sinusoidal function. A similar analysis was used in Horner et al. (2016), with the exception that they used a contrast to look for brain regions in which the sinusoidal model fit significantly better for one condition versus another (i.e., imagined navigation versus stationary periods).

#### 2.6.2. Comparing activity associated with aligned versus misaligned events

It is possible also to compare parameter estimates associated with aligned versus misaligned grid events. For example, in a subsequent analysis, Doeller et al. (2010) separated grid events into two regressors comprising those translations aligned within 15^o^ of a grid axis versus those more than 15^o^ from a grid axis, and again showed that significantly greater activity in entorhinal cortex was associated with events aligned with grid axes. This analysis strategy was used also by Kunz et al. (2015) who found that participants at increased genetic risk of Alzheimer’s disease show reduced BOLD response, relative to control participants, when contrasting trials ‘aligned > misaligned’ with the grid axis (i.e., a reduction in the ability to detect the grid code). Constantinescu et al. (2016) used a variation of the aligned versus misaligned analysis by sorting the grid event angles into 12 different regressors, each representing a 30^o^ bin. Six regressors comprised aligned trials, those events within ±15^o^ of the mean grid orientation (or a 60^o^ multiple of it). The remaining six regressors comprised misaligned trials, that is events offset from the mean grid orientation by 30^o^ (plus a 60^o^ multiple of this value) ±15^o^, and parameter estimates were extracted for each regressor.

#### 2.6.3. Analysis of grid code stability

The ability to detect the grid code in fMRI can be affected by the stability of the estimated grid orientation either between voxels within an ROI, or within voxels across different scanning runs and/or conditions (e.g., stability over time or different spatial environments, respectively). In terms of grid orientation stability between voxels within an ROI, if all voxels provide a different orientation value, then the resulting mean grid orientation would be random, and the coding of grid events in the test data depending on their deviation from the mean grid orientation would be arbitrary. To test whether there was evidence of coherence in the orientation of the grid code between different voxels in their entorhinal cortex ROI, Doeller et al. (2010) submitted all voxel orientation values to Rayleigh’s test for non-uniformity of circular data. Doeller et al. reported significant clustering of estimated orientations in around three-quarters of their participants.

Alternatively, an inability to detect grid codes in the fMRI signal could result from instability of the estimated grid orientation within a voxel over time. Kunz et al. (2015) tested the stability of the grid orientation over time by extracting the orientation of a voxel in one half of the data and comparing this to the same voxel’s orientation in the second half of the data. These data were scored such that if the values were within ±15^o^ of one another, then the grid orientation for the voxel was classified as stable. At the ROI level, the percentage of voxels showing stability in their estimated orientation over time could then be calculated. Even though participants at risk of Alzheimer’s disease showed coherence in grid orientation between voxels within a single scanning run, over time the orientation estimates for a given voxel differed. It was concluded, therefore, that the reduced ability to detect grid codes in the risk group resulted from instability in the orientation within-, but not between-, voxels in the Alzheimer’s risk group.

#### 2.6.4. Control analyses

Given that grid cells identified in rodents show a strict six-fold symmetry in their firing, it is necessary to test whether the best fit for the grid code analysis in fMRI is also a six-fold model, or whether other sinusoidal models fit the data equally well. In all studies published to date, the six-fold model has proven a better fit to estimate the orientation of the grid code in comparison to other symmetrical models (three-, four-, five-, seven- and eight-fold models; Horner et al., 2016; Kunz et al., 2015; Constantinescu et al., 2016; Doeller et al., 2010). These different models can be implemented in GridCAT, allowing the user to examine whether the six-fold model provides a better fit to the data.

An alternative control analysis, which can be carried out using the GridCAT, is to test for the grid code in regions where one would not expect to observe this signal (e.g., the visual cortex). Although this type of control analysis has been used previously (Doeller et al., 2010), it may be difficult to predict exactly which regions one would expect to see this pattern of data. For example, using an orthogonal localiser contrast, Constantinescu et al. (2016) found evidence of the sinusoidal response in a number of different regions including the ventromedial prefrontal cortex, and the posterior cingulate cortex.

## 3. Analysis of example dataset

To demonstrate some of the key features of the GridCAT, we detail here the analysis of functional data from an example participant who was scanned whilst they completed a spatial navigation task. The complete dataset of this example participant is available for download, so that the complete analysis pipeline described here can be reproduced using the GridCAT, giving the user the opportunity to explore its tools and functions. Futhermore, all necessary steps to analyse the example dataset are described in detail in the GridCAT manual.

### 3.1. Methods

The example participant was 28 years old, right handed, had normal vision and no history of psychiatric or neurological disorders.

#### 3.1.1. Spatial navigation task

Prior to scanning, the participant was asked to navigate a square virtual room (160 × 160 virtual meters) using a joystick and learn the location of three target-objects. Afterwards, the participant underwent two separate runs of fMRI scanning during which the participant navigated in the same virtual room. Each trial had the following structure: At the start, all target-objects disappeared and an image of one of them was shown at the bottom of the screen (Figure 2). The participant was asked to navigate to the position of the cued target-object and confirm their choice of location with a button-press. After the button-press, feedback was given to the participant via the target-object appearing at its correct location and a smiley-face displayed on the screen that was either green (if the ‘error distance’ between the correct location and the participant’s response was below 20 virtual meters), yellow (for ‘error distances’ between 20 and 30 virtual meters), or red (for ‘error distances’ larger than 30 virtual meters). Each scanning run lasted 16 minutes, and the participant was asked to complete as many trials as possible.

**Figure 2.**
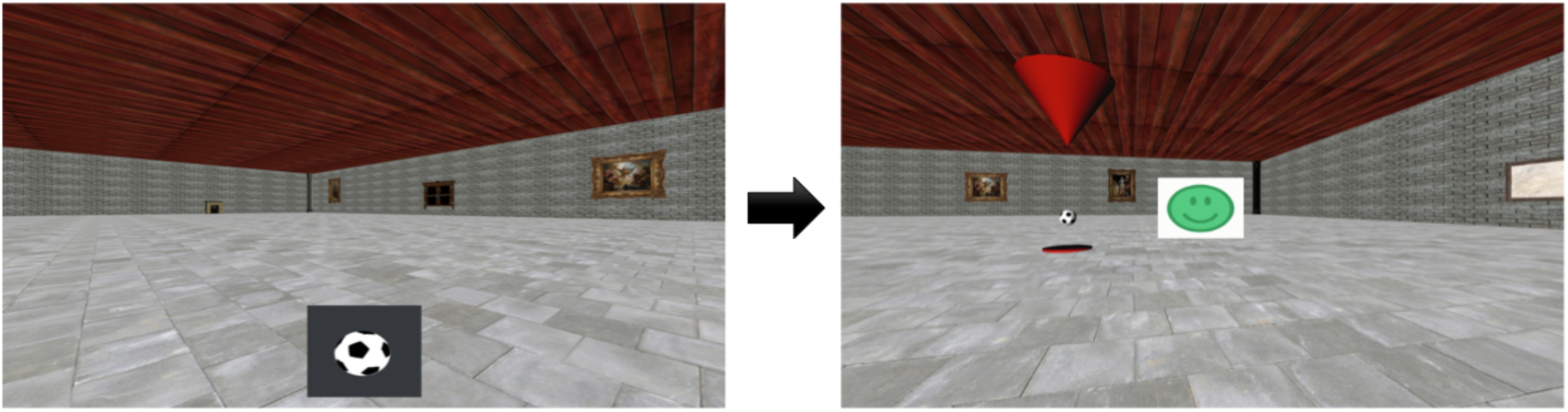
Example trial during fMRI scanning. Prior to scanning, the participant learned the locations of three target-objects in a virtual environment. Left: During scanning, one of the target-objects was cued (e.g., a football) and the participant was asked to navigate to its location. Right: After pressing a button to confirm their choice of location, the target-object appeared at its correct location and a smiley-face provided feedback as to the accuracy of the response.

#### 3.1.2. Scanning parameters

T2*-weighted functional images were acquired on a 3T Siemens Magnetom Prisma scanner using a partial-volume echo-planar imaging (EPI) sequence with the following parameters: repetition time (TR) = 1500 ms, echo time (TE) = 30 ms, slice thickness = 2 mm, in-plane-resolution = 2 × 2 mm, number of slices = 24, field of view = 216 mm, flip angle = 80°, slice acquisition order = interleaved.

For manual delineation of the entorhinal cortex, a high-resolution T2-weighted structural image was acquired using a turbo-spin-echo (TSE) sequence with the following parameters: repetition time (TR) = 6000 ms, echo time (TE) = 71 ms, slice thickness = 2 mm, in-plane-resolution = 0.5 × 0.5 mm, number of slices = 64, field of view = 224 mm, flip angle = 120°, slice acquisition order = interleaved.

#### 3.1.3. Analysis pipeline

Prior to analyses using the GridCAT, the functional images for the two runs were realigned and smoothed (5mm FWHM) using SPM12 (http://www.fil.ion.ucl.ac.uk/spm/). Anatomical masks of the left and right entorhinal cortices were traced manually (following Ding et al., 2016) on the participant’s T2-weighted image using ITK-SNAP (http://www.itksnap.org/), and co-registered to the EPI data. These two anatomical masks were used as separate ROIs for all following analyses.

As detailed in Section 2, there are a number of different ways grid codes can be examined in fMRI data, which are available to the GridCAT user. It is beyond the scope of this paper to demonstrate all possible combinations of modelling options; therefore, we chose a subset of parameters for the grid code analysis detailed here. The first parameter relates to the way in which the mean grid orientation is calculated. In GLM1, the GridCAT generates an image containing voxel-wise grid orientations, which can then be used to determine the mean grid orientation for a given ROI. The mean grid orientation can be calculated by averaging over voxels in the ROI either within individual scanning runs, or across multiple runs. For example, if one predicts that the grid orientation will change over runs, perhaps due to an experimental manipulation that could induce grid cell remapping (Fyhn et al., 2007), it would be sensible to estimate the grid orientation within individual runs, rather than averaging across them. Although we did not predict that there would be any changes in grid orientation over the two runs in our paradigm, we demonstrate the effect of estimating the mean grid orientation within versus across runs.

We examined also two different ways in which the grid events (i.e., translational movements within the virtual environment) can be modelled in GLM2. In one model, grid events were modelled using a single parametric modulator regressor (e.g., Doeller et al., 2010). In an alternate model, replicating the analysis of Kuntz et al. (2015), grid events were separated into two regressors - aligned or misaligned to the mean grid orientation - and contrasted with one another (‘aligned > misaligned’). The approach used by Constantinescu et al. (2016) in which grid events are separated into 12 different regressors comprising 30^o^ bins was not used here because our paradigm allowed for free exploration of the environment and therefore it is possible that not all directions were sampled equally. In all GLMs, we included as regressors of no interest the feedback phase in the paradigm, head motion parameters (*x*, *y*, *z*, yaw, pitch, and roll) derived from realignment in SPM, and the unused grid events (i.e., the grid events for GLM2 when fitting GLM1, and vice-versa).

Finally, we show how different symmetrical models (four-, five-, six-, seven-, and eight-fold) affect the model fit, with the prediction that the six-fold symmetrical model should provide the highest parameter estimates, given that this reflects grid cell firing symmetry.

### 3.2. Results

Consistent with the analysis strategy of Doeller et al. (2010), in GLM1 we found that the orientations of grid codes in voxels of both left and right entorhinal cortex showed significant non-uniformity, or clustering (see Figure 3). The GridCAT produces polar histogram plots, which indicate the different orientations derived from voxels in a given ROI, and the number of voxels sharing similar orientations. In these interactive plots, the mean grid orientation of all voxels within the ROI can also be calculated and plotted by the GridCAT. Moreover, Rayleigh’s test for non-uniformity of circular data can be carried out (applying code from the open-source toolbox CircStat2012a; Berens, 2009), in order to test whether the orientations of the grid code in voxels within an ROI show greater clustering than would be expected by chance. The example data suggest, therefore, that there is stability in grid orientation between voxels within the entorhinal cortex. As can be seen in Figure 3, the voxel-wise orientations estimated in the two separate runs were similar to one another, suggesting that the mean grid orientation could be calculated across both runs and used to categorize grid events in GLM2. If, however, these plots had indicated that the mean grid orientations changed over runs, the user might consider estimating and testing grid orientations within individual runs so that the categorization of grid events in GLM2, according to their alignment with the mean grid orientation, was more accurate. Furthermore, the GridCAT allows for the export of voxel-wise orientation values within an ROI, in order for additional analyses and/or statistical tests to be conducted on these data, depending on the user’s specific research question.

**Figure 3.**
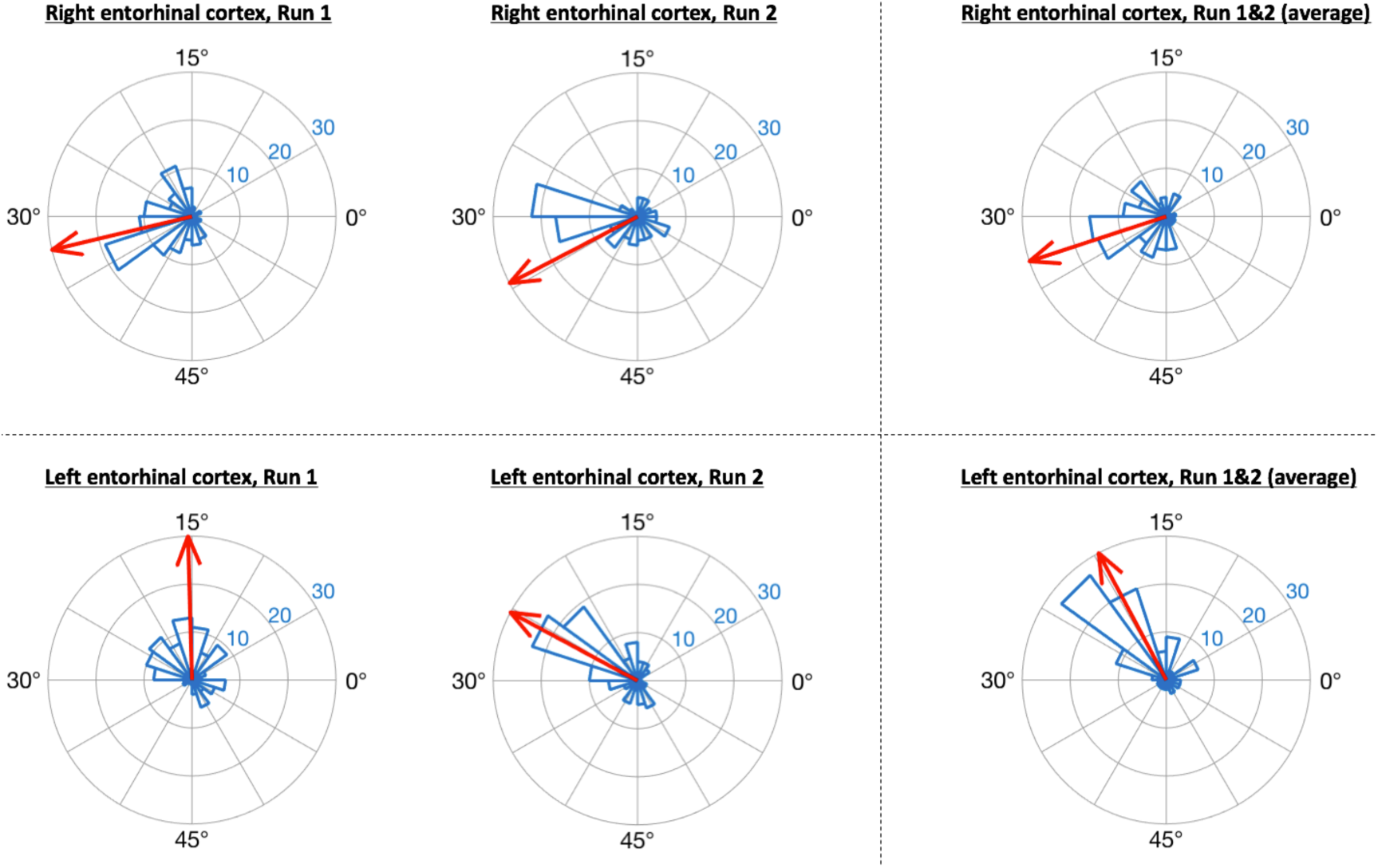
GridCAT polar histogram plots showing coherence of the grid orientation between-voxels in right and left entorhinal cortex ROIs. The length of each bar indicates the number of voxels that share a similar grid orientation, and the blue numbers indicate the number of voxels represented by each ring of the polar plot. The GridCAT also allows the user to calculate and visualize the mean grid orientation (red arrow) for each plot (which is used in GLM2 to model grid events with respect to their deviation from the mean grid orientation). Depending on the user‘s choice of model, the mean grid orientation can be calculated separately for run 1 (left column) and run 2 (middle column), or alternatively, the mean grid orientation over multiple runs (right column) can be calculated by averaging the parameter estimates. Furthermore, users can choose to carry out Rayleigh’s test for non-uniformity of circular data. For the present data, Rayleigh‘s test indicated that voxels in both the right (top row) and left (bottom row) entorhinal cortex showed significant clustering (i.e., coherence) in their orientations (all p < .00001).

The GridCAT can test also the within-voxel stability of the grid orientation across different scanning runs and/or conditions. When the user inputs two different voxel-wise orientation images derived from GLM1, and an ROI, the toolbox generates a plot comprising two polar plot rings (see Figure 4). For the analysis presented here, each ring represents a different scanning run, and circle markers denote the grid orientation of individual voxels; straight lines connect grid orientations of the same voxel across different runs. By default, the orientation of the grid code in a voxel is considered stable if the two values are within ±15^o^ of one another (i.e., the same threshold used in Kunz et al., 2015), and the GridCAT outputs the proportion of voxels within an ROI surviving this threshold. The stability of individual voxels is also displayed via the color of the connecting line; here, the GridCAT has displayed stable voxels in green and unstable voxels in red. Consistent with Kunz et al. (2015), the grid orientation for the example participant was consistent across the two runs, such that 75% of voxels in the right entorhinal cortex, and 60% of voxels in the left entorhinal cortex, maintained a stable orientation. The GridCAT provides the user with several other options in an interactive plot, including the ability to change the default value for threshold for stability (i.e., ±15^o^) if the researcher wishes to be more conservative or liberal with this estimate. Moreover, the user can specify several aesthetics of the plot, such as the colors and styles of the lines and markers.

**Figure 4.**
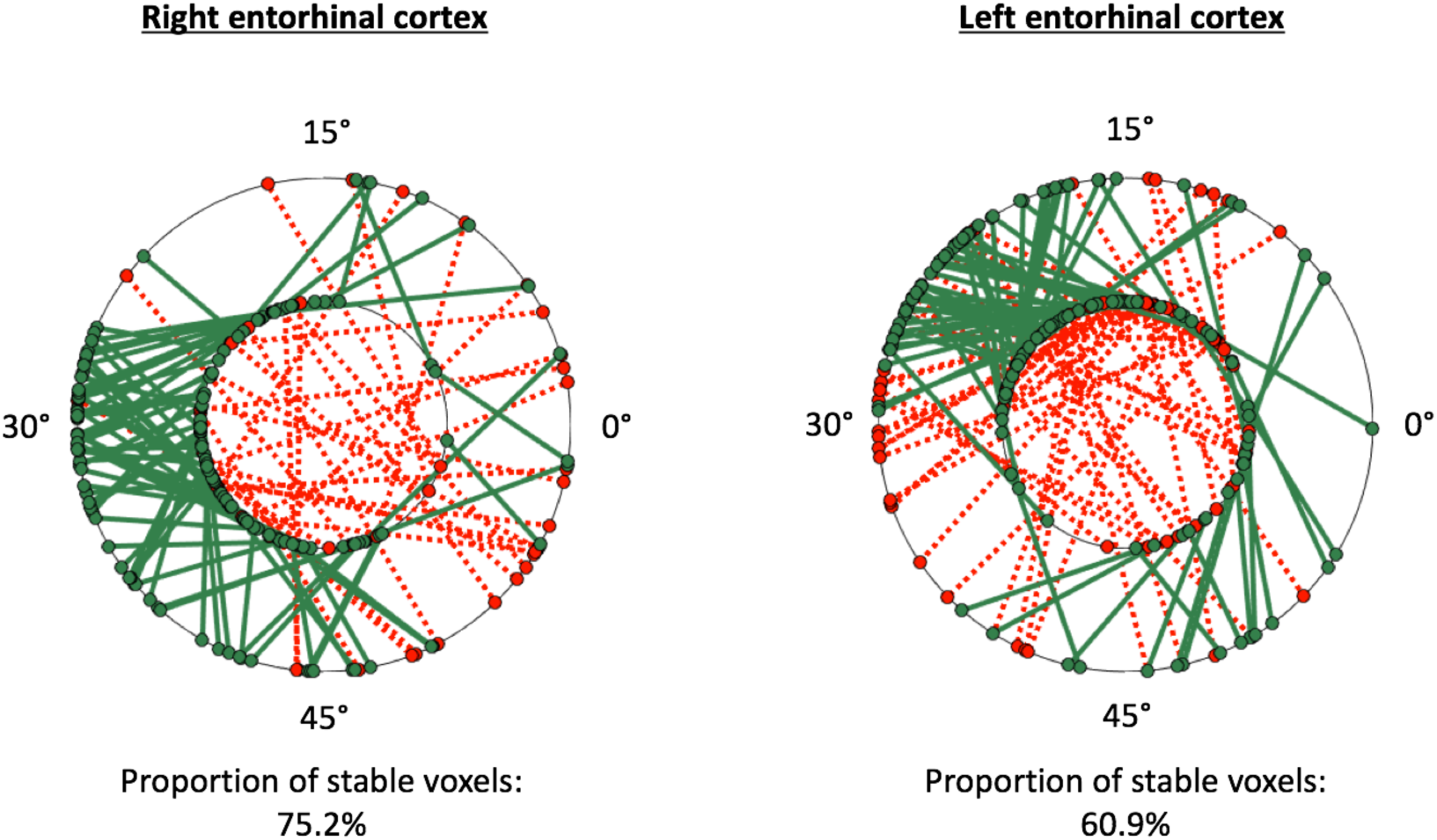
GridCAT polar plots showing coherence of the grid orientation within-voxels, across runs 1 and 2, in right and left entorhinal cortex ROIs. In both the right and left entohrinal cortex, the majority of voxels maintained the same grid orientation (±15°) across the two runs; the proportion of voxels maintaining the same orientation across runs is calculated automatically for the user. The two black rings in each plot represent the two different runs (inner ring: run 1, outer ring: run 2), and the orientation of the grid code for each voxel is indicated with a circular marker; a line connects the orientations of each voxel. Green solid lines indicate voxels with stable orientations, whereas red dotted lines indicate voxels with unstable orientations. The GridCAT allows the user also to customize the plots, including the color schemes, line styles, as well as adapting the threshold for classifying a voxel as stable.

For the test data in GLM2, the GridCAT allows users to model grid events either with a parametric modulator regressor (e.g., Doeller et al., 2010), or by separating grid events into trials aligned versus misaligned with the mean grid orientation and contrasting these values (‘aligned > misaligned’ see Sections 2.6.1 and 2.6.2 for more details). The two methods resulted in comparable parameter estimates in the right entorhinal cortex ROI, with the ‘aligned > misaligned’ contrast method associated with slightly higher parameter estimates relative to the parametric modulator (see Figure 5). In the left entorhinal cortex, there were less obvious differences between methods, however the ‘aligned > misaligned’ contrast again yielded the highest parameter estimate, but only when the mean average grid orientation was calculated using the data from both runs in GLM1. That the grid code metrics appear generally stronger in the right hemisphere, in terms of between-voxel and within-voxel grid orientation coherence, and model fit in GLM2, supports previous findings (Doeller et al., 2010). It is unclear from a theoretical viewpoint, however, why this should be the case, and requires more extensive comparisons within individual subjects to determine the consistency of this effect.

The control analysis tests for the fit of different symmetrical models, to examine whether the six-fold symmetry explains best the data. Consistent with the other results reported here, in right entorhinal cortex the six-fold symmetrical model resulted in the numerically highest parameter estimates relative to all other models. In the left entorhinal cortex, the six-, seven-, and eight-fold models all appear to fit the data equally well (Figure 5). It should be noted, however, that other papers reporting a better fit of the six-fold symmetrical model show effects at the group-level, rather than within individual subjects. Accordingly, there may be substantial variability in these estimates both inter-subject, as well as intra-subject, as demonstrated here by the difference between left and right hemispheres.

**Figure 5.**
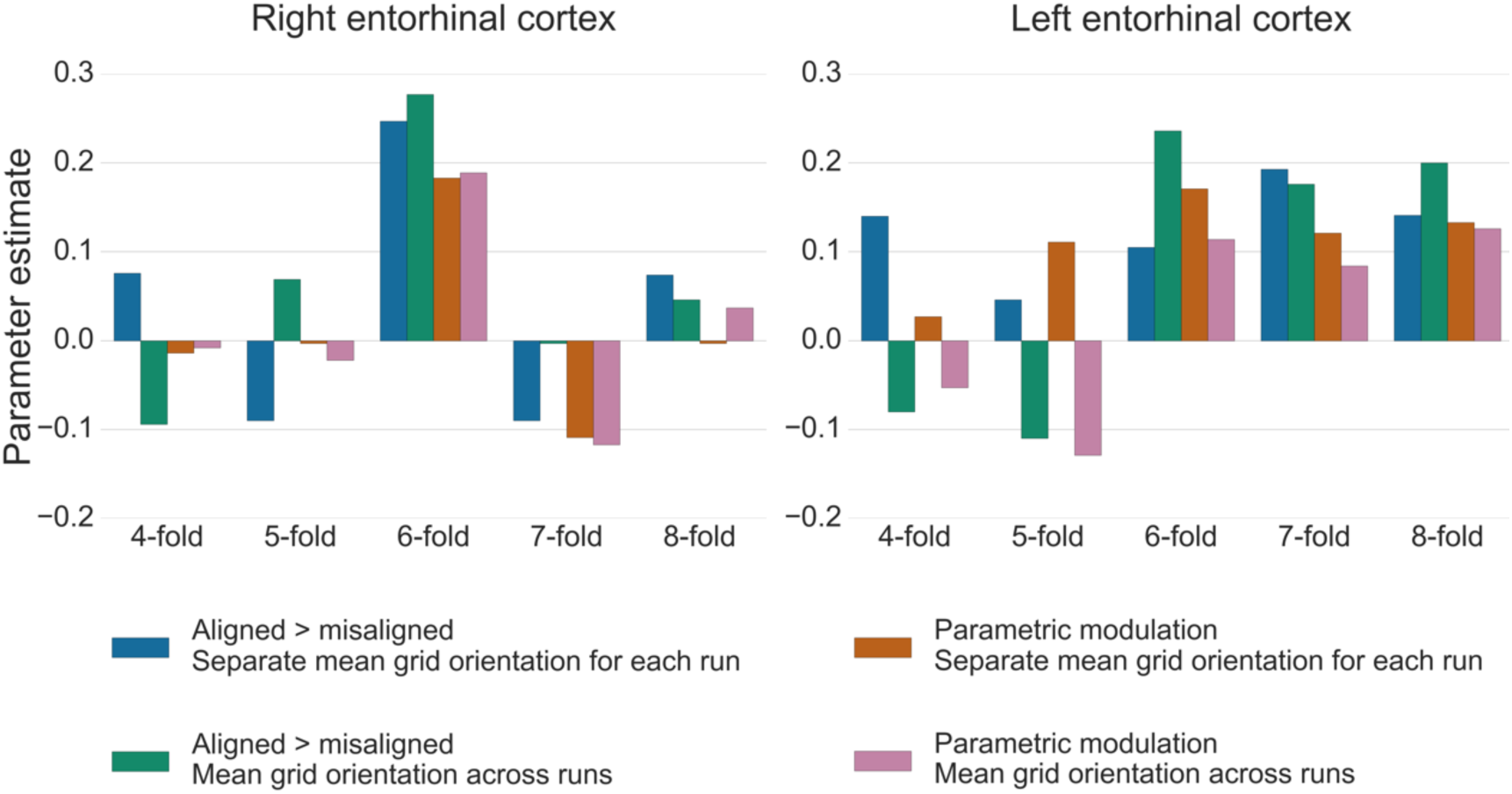
Parameter estimates associated with grid events using different model designs and grid code symmetries. In the right entorhinal cortex, the six-fold symmetrical provided the best model fit, and the ‘aligned > misaligned’ contrast yielded numerically higher parameter estimates relative to the single parametric modulator regressor contrast; calculating the mean grid orientation over two runs, versus using separate mean grid orientations for each run, did not impact upon model fit. In the left entorhinal cortex, the ‘aligned > misaligned’ contrast again provided the highest parameter estimate, however the effect of generating the mean grid orientation over multiple runs versus separate single runs was more variable. Moreover, other symmetrical models provided equally good model fits for this participant. All bars show the mean parameter estimate averaged over all voxels within right and left entorhinal cortex, respectively.

## 4. Discussion

The GridCAT is an open source toolbox allowing researchers to examine the putative firing of grid cells (i.e., grid codes) in human fMRI data. The GridCAT provides a simple GUI, and accompanying open-source code, for the analysis of fMRI data, so that the user can conduct the entire grid code analysis pipeline. In order to learn and understand the functionality of the GridCAT, a detailed manual is provided to guide the user through all analysis steps, and the user can also follow the instructions to analyse an example dataset. Furthermore, example scripts are provided for those who do not want to use the GUI, but rather use and modify the existing open-source code of the GridCAT.

Despite the great deal of research into grid cells using non-human animal species (for an overview see Rowland et al., 2016), there remain very few studies examining grid codes in human fMRI. Given that this cellular mechanism is now purported to support more than just pure spatial navigation behaviour in humans (Constantinescu et al., 2016), researchers now face the exciting challenge of elucidating exactly what role this cell may play in other cognitive domains. One potential reason why there remain so few papers examining grid codes in humans may be that the analysis method is distinctly non-trivial, requiring knowledge not only of fMRI data analysis but also of other domains, such as computer programming and quadrature filter analysis. The GridCAT opens up this cutting-edge research area to researchers less comfortable with programming by allowing users to analyse data using a GUI; researchers confident with programming, however, can edit the open-source code accompanying the GridCAT to implement their own analysis pipelines. Importantly, the toolbox requires only a licence for Matlab and the freely available SPM12 toolbox (http://www.fil.ion.ucl.ac.uk/spm/). Moreover, fMRI preprocessing can be carried out in the researcher’s neuroimaging analysis package of choice.

The toolbox is designed to be flexible in allowing the user to implement the analysis pipelines that have been used previously in grid code publications. These include different ways in which the data can be partitioned for GLM1 and GLM2, different ways to model aligned versus misaligned trials, and the calculation of a variety of grid code metrics to assess grid code stability and symmetrical model fit. As noted above, although it is beyond the scope of this paper to compare the results of all different model selection parameters, we believe that this is an important goal for the field so that researchers will have a better idea as to the factors that aid detection of these signals in fMRI data. By making all of these options available to the user, and the wider neuroscience community, the GridCAT has provided the first step in achieving this goal and has the potential to accelerate grid code research in humans.

## Acknowledgements

This work was supported by the European Research Council Starting Investigator Grant AGESPACE (335090) and by the Collaborative Research in Computational Neuroscience Grant (01GQ1303) of the German Ministry of Education and Research (BMBF).

## References

Barry, C., Hayman, R., Burgess, N., & Jeffery, K. J. (2007). Experience-dependent rescaling of entorhinal grids. Nature Neuroscience, 10(6), 682–684. http://doi.org/10.1038/nn1905

Bellmund, J. L., Deuker, L., Navarro Schröder, T., & Doeller, C. F. (2016). Grid-cell representations in mental simulation. eLife, 5, 12897–12901. http://doi.org/10.7554/eLife.17089

Berens, P. (2009). CircStat: A MATLAB Toolbox for Circular Statistics. Journal of Statistical Software, 31(10), 1–21. http://doi.org/10.7554/eLife.17089

Constantinescu, A. O., O’Reilly, J. X., & Behrens, T. E. J. (2016). Organizing conceptual knowledge in humans with a gridlike code. Science, 352(6292), 1464–1468. http://doi.org/10.1126/science.aaf0941

Ding, S.-L., Royall, J. J., Sunkin, S. M., Ng, L., Facer, B. A. C., Lesnar, P., Guillozet-Bongaarts, A., McMurray, B., Szafer, A., Dolbeare, T. A., Stevens, A., Tirrell, L., Benner, T., Caldejon, S., Dalley, R. A., Dee, N., Lau, C., Nyhus, J., Reding, M., Riley, Z. L., Sandman, D., Shen, E., Kouwe, van der A., Varjabedian, A., Write, M., Zollei, L., Dang, C., Knowles, J. A., Koch, C., Phillips, J. W., Sestan, N., Wohnoutka, P., Zielke, H. R., Hohmann, J. G., Jones, A. R., Bernard, A., Hawrylycz, M. J., Hof, P. R., Fischl, B., & Lein, E. S. (2016). Comprehensive cellular-resolution atlas of the adult human brain. Journal of Comparative Neurology, 524(16), 3127–3481. http://doi.org/10.1002/cne.24080

Doeller, C. F., Barry, C., & Burgess, N. (2010). Evidence for grid cells in a human memory network. Nature, 463(7281), 657–661. http://doi.org/10.1038/nature08704

Fyhn, M., Molden, S., Witter, M. P., Moser, E. I., & Moser, M.-B. (2004). Spatial Representation in the Entorhinal Cortex. Science, 305(5688), 1258–1264. https://doi.org/10.1126/science.1099901

Fyhn, M., Hafting, T., Treves, A., Moser, M.-B., & Moser, E. I. (2007). Hippocampal remapping and grid realignment in entorhinal cortex. Nature, 446(7132), 190–194. http://doi.org/10.1038/nature05601

Hafting, T., Fyhn, M., Molden, S., Moser, M.-B., & Moser, E. I. (2005). Microstructure of a spatial map in the entorhinal cortex. Nature, 436(7052), 801–806. http://doi.org/10.1038/nature03721

Horner, A. J., Bisby, J. A., Zotow, E., Bush, D., & Burgess, N. (2016). Grid-like Processing of Imagined Navigation. Current Biology, 26(6), 842–7. http://doi.org/10.1016/j.cub.2016.01.042

Jacobs, J., Weidemann, C. T., Miller, J. F., Solway, A., Burke, J. F., Wei, X.- X., Suthana, N., Sperling, M. R., Sharan, A. D., Fried, I., & Kahana, M. J. (2013). Direct recordings of grid-like neuronal activity in human spatial navigation. Nature Neuroscience, 16(9), 1188–1190. http://doi.org/10.1038/nn.3466

Killian, N. J., Jutras, M. J., & Buffalo, E. A. (2012). A map of visual space in the primate entorhinal cortex. Nature, 491(7426), 761–4. https://doi.org/10.1038/nature11587

Kunz, L., Schröder, T. N., Lee, H., Montag, C., Lachmann, B., Sariyska, R., Reuter, M., Stirnberg, R., Stöcker, T., Messing-Floeter, P. C., Fell, J., Doeller, C. F., & Axmacher, N. (2015). Reduced grid-cell-like representations in adults at genetic risk for Alzheimer’s disease. Science, 350(6259), 430–433. http://doi.org/10.1126/science.aac8128

Nili, H., Wingfield, C., Walther, A., Su, L., Marslen-Wilson, W., & Kriegeskorte, N. (2014). A Toolbox for Representational Similarity Analysis. PLoS Computational Biology, 10(4), e1003553. http://doi.org/10.1371/journal.pcbi.1003553

O’Keefe, J., & Dostrovsky, J. (1971). The hippocampus as a spatial map. Preliminary evidence from unit activity in the freely-moving rat. Brain Research, 34(1), 171–175. http://doi.org/10.1016/0006-8993(71)90358-1

Oosterhof, N. N., Connolly, A. C., & Haxby, J. V. (2016). CoSMoMVPA: Multi-Modal Multivariate Pattern Analysis of Neuroimaging Data in Matlab/GNU Octave. Frontiers in Neuroinformatics, 10, 27. http://doi.org/10.3389/fninf.2016.00027

Rowland, D. C., Roudi, Y., Moser, M.-B., & Moser, E. I. (2016). Ten Years of Grid Cells. Annual Review of Neuroscience, 39(1), 19–40. http://doi.org/10.1146/annurev-neuro-070815-013824

Sargolini, F., Fyhn, M., Hafting, T., McNaughton, B. L., Witter, M. P., Moser, M.-B., & Moser, E. I. (2006). Conjunctive representation of position, direction, and velocity in entorhinal cortex. Science, 312(5774), 758–62. http://doi.org/10.1126/science.1125572

Yartsev, M. M., Witter, M. P., & Ulanovsky, N. (2011). Grid cells without theta oscillations in the entorhinal cortex of bats. Nature, 479(7371), 103–107. https://doi.org/10.1038/nature10583

